# Autism spectrum disorder: understanding the impact of SNPs on biological pathways in the fetal and adult cortex

**DOI:** 10.1101/2021.03.03.433667

**Authors:** E. Golovina, T. Fadason, T.J. Lints, C. Walker, M.H. Vickers, J.M. O’Sullivan

**Affiliations:** Liggins Institute, University of Auckland; Maurice Wilkins Centre, University of Auckland, Auckland, New Zealand; School of Medical Science, University of Auckland; School of Population Health, University of Auckland; Brain Research New Zealand, University of Auckland, Auckland, New Zealand; MRC Lifecourse Epidemiology Unit, University of Southampton, United Kingdom

## Abstract

Autism spectrum disorder (ASD) is a neurodevelopmental disorder characterized by significant and complex genetic etiology. GWAS studies have identified genetic variants associated with ASD, but the functional impacts of these variants remain unknown. Here, we integrated four distinct levels of biological information (GWAS, eQTL, spatial genome organization and protein-protein interactions) to identify potential regulatory impacts of ASD-associated SNPs (*p* < 5×10^-8^) on biological pathways within fetal and adult cortical tissues. We found 80 and 58 SNPs that mark regulatory regions (i.e. expression quantitative trait loci or eQTLs) in the fetal and adult cortex, respectively. These eQTLs were also linked to other psychiatric disorders (e.g. schizophrenia, ADHD, bipolar disorder). Functional annotation of ASD-associated eQTLs revealed that they are involved in diverse regulatory processes. In particular, we found significant enrichment of eQTLs within regions repressed by Polycomb proteins in the fetal cortex compared to the adult cortex. Furthermore, we constructed fetal and adult cortex-specific protein-protein interaction networks and identified that ASD-associated regulatory SNPs impact on immune pathways, fatty acid metabolism, ribosome biogenesis, aminoacyl-tRNA biosynthesis and spliceosome in the fetal cortex. By contrast in the adult cortex, they largely affect immune pathways. Overall, our findings highlight potential regulatory mechanisms and pathways important for the etiology of ASD in early brain development and adulthood. This approach, in combination with clinical studies on ASD, will contribute to individualized mechanistic understanding of ASD development.

## Introduction

Autism spectrum disorder (ASD) represents a heterogeneous group of closely related conditions that are characterized by early-appearing social communication deficits and restricted, repetitive or unusual sensory-motor behaviours [1]. Epidemiological studies estimate that approximately 1% of people worldwide have ASD [2].

Over the past decade, genome-wide association (GWAS) and genetic studies have identified increasing numbers of single nucleotide polymorphisms (SNPs) [3, 4] and other forms of variation (e.g., copy number variants, rare structural variants) [5, 6] that are associated with ASD. The proportion of ASD explained by SNPs has been estimated to be between 17-60% [7, 8], thus their contribution should not be neglected. However, the functions of the genetic variants that are responsible for the association with ASD remain poorly characterized. As such, we do not yet fully understand how to translate information on ASD-associated SNPs into specific biological mechanisms that can be therapeutically targeted to alleviate the symptoms and complications of ASD.

The majority of ASD-associated SNPs are located within the non-coding components of the genome. Studies of non-coding disease-associated SNPs have demonstrated that they can mark regulatory elements that alter gene expression [9, 10]. Notably, these regulatory elements are only associated with the expression (eQTL or expression quantitative trait locus) of the adjacent gene in ∼40% of cases [11]. The remaining 60% of the identified eQTLs involve interactions with non-adjacent genes that can be >1Mb away in the linear DNA sequence or even on a different chromosome. The regulatory effects can occur in trans (e.g. miRNA) or by spatial associations of the regulatory element and target gene. As such, the three-dimensional (3D) genome organization, which emerges from the sum of the biophysical interactions within the nucleus, includes tissue-specific spatial interactions between eQTL regions and the genes that they control (hereafter eGenes) [12]. These spatial interactions are dynamic, developmentally and temporally dependent [13]. Thus, integrating biological measurements on developmental and tissue-specific spatial chromatin interactions with eQTL information could inform our understanding of the regulatory impacts of ASD-associated SNPs.

ASD is widely considered to be a neurodevelopmental disorder resulting from functional changes within the brain. There are studies connecting cortical dysfunctions and ASD using imaging [14], cortical architecture [15], or gene expression [16]. Therefore, characterizing the functional impacts (*i.e.* on gene regulation) of the ASD-associated SNPs and translating them to the affected biological pathways in fetal and adult cortical tissues may provide mechanistic insights into the etiology of ASD during neurodevelopment.

Using protein-protein interaction (PPI) networks to explore interactions between proteins encoded by known disease-associated genes is a powerful approach to study the etiology of complex diseases, including psychiatric disorders [17, 18]. PPI network analyses have been used to discover essential proteins, clusters of proteins with similar, overlapping or combinatorial functions, and associated pathways involved in tissue-specific contributions to ASD etiology [17, 18]. However, the potential contributions of cortex-specific developmental changes to these networks in ASD development have yet to be investigated.

Here, we integrated ASD-associated GWAS SNPs with cortex-specific 3D genome structure and eQTL information to identify genes that are spatially regulated in fetal (14-21 postconceptional weeks) and adult (21-70 years of age) cortical tissues. We incorporated cortex-specific expression patterns and PPI networks to identify candidate genes and pathways that have putative roles in the etiology of ASD-associated changes in the cortex. The identified gene sets were enriched for immune pathways, fatty acid metabolism, ribosome biogenesis, aminoacyl-tRNA biosynthesis and the spliceosome in the fetal cortex. By contrast, the adult cortical gene set was largely related to immune pathways. Collectively, our results provide insight into potential cortex-specific regulatory mechanisms and pathways through which ASD-associated SNPs can contribute to the development and maintenance of ASD.

## Methods

### Data access approvals

Data access was approved by the dbGaP (https://www.ncbi.nlm.nih.gov/gap/) Data Access Committee(s) for: 1) cortical plate and germinal zone neuron Hi-C datasets (project #16489: “Finessing predictors of cognitive development”, accession: phs001190.v1.p1) [19]; 2) total RNA-seq and WGS datasets across GTEx v8 tissues (project #22937: “Untangling the genetics of disease multimorbidity”, accession: phs000424.v8.p2) [20]; and 3) total RNA-seq and genotyping datasets for fetal brain cortical tissue from 14-21 postconceptional weeks (PCWs) (project #25321: “Gene regulatory networks in Autism”, accession: phs001900.v1.p1) [21] (Supplementary Table 1).

### GTEx data processing

Genotypes (derived by Whole Genome Sequencing) were processed by the Genotype-Tissue Expression (GTEx) project, and filtered genotypes (with minor allele frequency ≥0.1) for 838 tissue donors were downloaded from dbGaP (https://www.ncbi.nlm.nih.gov/gap/, 01/05/2020). RNA-seq data were processed by GTEx using RNA-seq analysis (https://github.com/broadinstitute/gtex-pipeline/tree/master/rnaseq) and eQTL discovery (https://github.com/broadinstitute/gtex-pipeline/tree/master/qtl) pipelines to calculate normalized gene expression and covariates. The resulting expression (GTEx_Analysis_v8_eQTL_expression_matrices.tar) and covariates (GTEx_Analysis_v8_eQTL_covariates.tar.gz) data were downloaded from GTEx website (https://www.gtexportal.org/home/datasets, 01/05/2020).

### Fetal RNA-seq data processing

Raw RNA-seq fastq files [21] were downloaded from dbGaP (05/06/2020), merged across lanes from the same sample (final dataset of 219 individuals) and analysed using FastQC (v0.11.9; default parameters). FastQC reports were visually inspected and there were no samples that did not pass the quality check (no failures for “Per base sequence quality”, “Per sequence quality scores”, “Per base N content” and “Sequence Length Distribution” metrics). All RNA-seq data were processed according to the GTEx pipeline (https://github.com/broadinstitute/gtex-pipeline/tree/master/rnaseq) (Supplementary Figure 1). The same reference genome and annotation files were used to calculate expression for fetal brain RNA-seq data. Briefly, merged fastq files were aligned to the GRCh38 reference genome (Homo_sapiens_assembly38_noALT_noHLA_noDecoy.fasta, gs://gtex-resources) using STAR (v2.5.3a). Duplicated mapped reads were marked using Picard MarkDuplicates module (v2.21.4). Quality control metrics and gene-level expression data were calculated using RNA-seQC (v2.3.6) on the basis of GENCODE v26 gene annotation (gencode.v26.GRCh38.genes.gtf, gs://gtex-resources). Sample-level gene read and TPM (Transcripts Per kilobase Million) counts were concatenated using combine_GCTs.py.

The GTEx eQTL discovery (https://github.com/broadinstitute/gtex-pipeline/tree/master/qtl) pipeline was further used to calculate normalized gene expression and covariates. Briefly, read counts were normalised using the TMM algorithm and genes were selected if they had counts of ≥0.1 TPM in In ≥20% samples and ≥6 unnormalized reads in ≥20% samples. Genes were inverse normal transformed across samples. Top 5 genotype principal components (calculated using compute_genotype_pcs.py script from https://github.com/broadinstitute/gtex-pipeline/tree/master/genotype), 30 PEER factors, sex and genotyping platform were used as covariate in the eQTL analysis.

### Fetal genotype data processing

Genotypes (derived by Array Genotyping) for 219 fetal brain donors [21] were downloaded from dbGaP (05/06/2020), processed and prepared in the GTEx format (Supplementary Figure 1). Briefly, data were preprocessed to correct strand orientation and position of the variants on the GRCh37 reference genome (update_build.sh script). Variants that do not have strand information for HumanOmni25-8v1-2_A1 and HumanOmni2-5Exome-8-v1-1-A genotyping chips were excluded (strand files and update_build.sh script were downloaded from https://www.well.ox.ac.uk/~wrayner/strand/, 01/07/2020, Supplementary Table 1).

Genotype data quality control was performed using PLINK (v2.0). Genetic variants were filtered based on Hardy-Weinberg equilibrium *p* <1×10^-6^, minor allele frequency 0.01 and variant missing genotype rate 0.05. Within-family IDs were used as sample IDs in the output vcf file (--recode vcf-iid bgz). In total, 663,956 variants passed QC filters. BCFtools (v1.10.2) was used to exclude genetic variants on chromosome 0 (omitted due to mapping to multiple locations) and chromosome 25 (XY pseudoautosomal region), to rename chromosomes 23 (X), 24 (Y) and 26 (MT), to fix REF allele, to check sample’s sex and to normalize the output vcf file to the GRCh37 reference genome (human_g1k_v37.fasta.gz, downloaded from ftp://ftp.1000genomes.ebi.ac.uk/vol1/ftp/technical/reference/, 01/07/2020). Normalized vcf files were further validated using VCFtools (v0.1.15).

The Sanger Imputation Service (https://imputation.sanger.ac.uk/, 02/07/2020) [22] was used to: 1) pre-phase the validated genotypes with Eagle (v2.4.1); and 2) to impute them using the 1000 Genomes Phase 3 multi-ethnic reference panel and PBWT algorithm. Imputation accuracy was assessed by filtering variants by info score (INFO<0.8). Imputed genotypes were filtered for Hardy-Weinberg equilibrium *p* <1×10^-6^, variant missing genotype rate 0.05 and minor allele frequency 0.01.

CrossMap (v0.2.6) was used to convert coordinates of genetic variants from genome build hg19 to hg38, resulting in ∼54.8 million genetic variants. BCFtools were used to set variant IDs according to the GTEx variant ID format (e.g. chr1_61170_C_T_b38 where chr1 is chromosome name, 61170 is variant position on the chromosome, C is reference allele, T is alternate allele and b38 is genome build 38). The resulting vcf file was converted to plink format and information on sample sex included. To create a lookup table, genetic variants were annotated with rsIDs from dbSNP build 151 database.

### eQTL mapping

Genotypes, expression matrices and covariates for fetal and adult brain were integrated into CoDeS3D [11] (https://github.com/Genome3d/codes3d-v2) pipeline as two separate eQTL databases Lastly, tensorQTL (https://github.com/broadinstitute/tensorqtl) algorithm was used to perform cis- and trans-QTL mapping.

### Hi-C data processing

In order to study spatial regulatory interactions in fetal and adult cortical tissues, we analysed two fetal brain-specific (i.e. cortical plate and germinal zone neurons) [19] and one adult cortex-specific (i.e. dorsolateral prefrontal cortex cells) [12] Hi-C chromatin interaction libraries (Supplementary Table 1). Raw Hi-C data were downloaded from dbGaP (accession: phs001190.v1.p1) and GEO (https://www.ncbi.nlm.nih.gov/geo/, accession: GSE87112) and analyzed using Juicer (v1.5) [23] (https://github.com/aidenlab/juicer) pipeline to generate Hi-C libraries. The pipeline included BWA (v0.7.15) alignment of paired-end reads onto the hg38 reference genome, merging paired-end read alignments and removing chimeric, unmapped and duplicated reads. The remaining read pairs we refer to as “contacts”. Only Hi-C libraries that contain >90% alignable unique read pairs, and >50% unique contacts (<40% duplication rate) within the total sequenced read pairs were included in the analysis. Files containing cleaned Hi-C contacts locations (*i.e.* *_merged_nodups.txt files) were processed to obtain Hi-C chromatin interaction libraries in the following format: read name, str1, chr1, pos1, frag1 mapq1, str2, chr2, pos2, frag2, mapq2 (str = strand, chr = chromosome, pos = position, frag = restriction site fragment, mapq = mapping quality score, 1 and 2 correspond to read ends in a pair). Reads where both ends had a mapq ≥30 were included in the final library. Hi-C chromatin interactions represent all captured pairs of interacting restriction fragments in the genome. As such, restriction fragments were used to identify regulatory interactions between SNPs and genes (Figure 1).

**Fig. 1.**
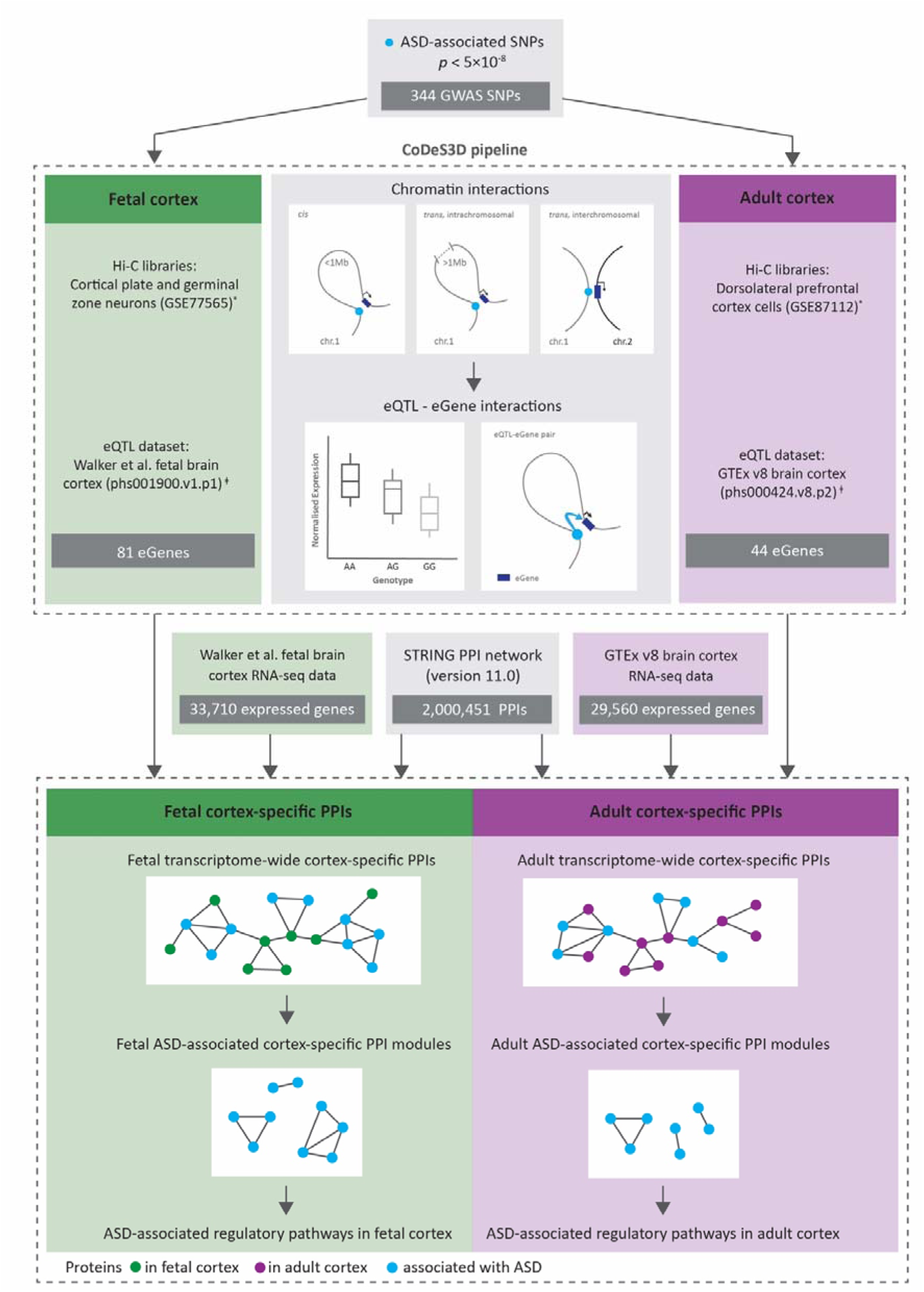
Overview of the analysis pipeline used in this study. 344 ASD-associated SNPs (*p*<5×10^-8^) represented in both fetal and adult cortex-specific eQTL datasets were run through the CoDeS3D pipeline to identify 81 and 44 spatially regulated genes in fetal and adult cortical tissues, correspondingly. Over 11 million protein-protein interactions (PPIs) were downloaded from STRING database (version 11.0) and combined with cortex-specific expression data (GTEx v8 or Walker et al. datasets) to construct tissue-specific transcriptome-wide PPI networks. Modules that were enriched with ASD-eQTL associated genes were identified in the fetal and adult cortical tissues. * Hi-C datasets for cortical plate and germinal zone neurons (phs001190.v1.p1) were obtained from Won et al [19], Hi-C datasets for adult dorsolateral prefrontal cortex cells were obtained from Schmitt et al [12]. □eQTL datasets for fetal and adult cortex were obtained from Walker et al [21] and GTEx v8 [20], correspondingly.

### Identification of SNPs associated with ASD

Single-nucleotide polymorphisms (SNPs) associated with ASD (n = 454) were downloaded from the GWAS Catalog (www.ebi.ac.uk/gwas/; 05/04/2020; Supplementary Table 2). Only SNPs associated with ASD with a *p* <5×10^-8^ were included in downstream analyses. Phenotypes were defined as the mapped traits associated with the SNP in the GWAS Catalog.

### Identification of spatial regulatory interactions using CoDeS3D

CoDeS3D [11] (https://github.com/Genome3d/codes3d-v2) was used to identify genes that spatially interact with putative regulatory regions tagged by ASD-associated SNPs (Figure 1, Supplementary Table 3). Briefly, the human genome reference (hg38) was fragmented at HindIII sites (A/AGCTT), the restriction enzyme that was used in the preparation of the Hi-C libraries. The CoDeS3D algorithm then checked if the ASD-associated SNP rsID numbers were present in the eQTL database and identified the restriction fragments that were tagged by the SNPs. Using fetal (i.e. cortical plate and germinal zone neurons) or adult (i.e. dorsolateral prefrontal cortex cells) cortex-specific Hi-C libraries, CoDeS3D identified the restriction fragments that were captured interacting with the SNP-tagged restriction fragments. Interacting fragments that overlapped annotated genes (GENCODE transcript model version 26) were identified. The resulting SNP-gene pairs were used to query adult cortex and fetal cortex eQTL databases to identify cis- and trans-acting eQTL-eGene interactions (*i.e.* genes, whose expression levels are associated with a SNP). Finally, significant cortex-specific eQTL-eGene interactions were identified using the Benjamini-Hochberg (BH) FDR correction to adjust the eQTL *p* values (FDR < 0.05) (Supplementary Table 3).

### Functional annotation of eQTL SNPs associated with ASD

The identified ASD-associated eQTLs were annotated using wANNOVAR tool [24] (http://wannovar.wglab.org/, 10/08/2020) to obtain information about the locus they tagged (Supplementary Table 4). Enrichment of the eQTLs within transcription factor binding sites was determined using SNP2TFBS (https://ccg.epfl.ch//snp2tfbs/, 07/09/2020) [25]. Enrichment within active regulatory elements and histone modification marks was identified using the Roadmap Epigenomics Project 15-state ChromHMM models [26, 27] for adult dorsolateral prefrontal cortex (E073_15_coreMarks_hg38lift_mnemonics.bed.gz) and fetal brain (E081_15_coreMarks_hg38lift_mnemonics.bed.gz) (downloaded from https://egg2.wustl.edu/roadmap/data/byFileType/chromhmmSegmentations/ChmmModels/coreMarks/jointModel/final/, 22/11/2020) (Supplementary Table 1). Enrichment analyses were performed using R package regioneR [28] (permutation test: 1000). Finally, we evaluated identified eQTL SNP associations with other phenotypes in the GWAS Catalog (downloaded on 26/08/2020) (Supplementary Table 4). Phenotypes were defined as the mapped traits associated with the SNP in the GWAS Catalog. Only SNP-phenotype associations with a *p*<5×10^-8^ were included in the analysis. All genomic positions and SNP annotations were obtained for human genome reference build hg38 (GRCh38) release 75.

### Construction of ASD-associated PPI networks

The STRING [29] PPI network (version 11.0, protein.links.full.v11.0.txt.gz, https://string-db.org/) was downloaded on 24/09/2020. We extracted 2,000,451 protein-protein interactions (with a combined score ≥400) between a total of 19,258 unique human proteins (Figure 1).

Transcriptome-wide fetal and adult cortex-specific PPIs (CSPPIs) were constructed by combining the STRING PPI network with cortex-specific expression data from GTEx v8 or fetal brain datasets (Figure 1). Ensembl protein (STRING) [29] and transcript identifiers (GTEx and Walker et al. RNA-seq data) were mapped to Ensembl gene identifiers. The CSPPIs represents subnetworks of the STRING PPI network, in which a protein/node is only present if it is expressed in the cortical tissue (adult or fetal). The size of each node depends on the protein expression levels (no missing values and minimum expression level >0 TPM) in the corresponding cortical tissue. Proteins that were not annotated in the expression datasets were also removed from the CSPPI network. Edges are only present if both interacting proteins are expressed in the cortical tissue. The resulting CSPPI networks contained 1,784,342 PPIs between 17,156 unique proteins in the adult brain, and 1,690,571 PPIs between 16,519 unique proteins in the fetal brain. To build ASD-specific fetal and adult CSPPIs, only interactions between ASD-associated genes we extracted from fetal and adult CSPPIs. The Louvain clustering algorithm [30] was further applied to identify ASD-specific clusters of functionally related proteins within the CSPPI networks.

### Gene Ontology enrichment and pathway analyses

Gene Ontology (GO) enrichment and pathways analyses for the eGenes within the ASD-specific CSPPI clusters were performed using the g:GOSt module of the g:Profiler tool [31]. eGene enrichment was tested within the biological process, molecular function and cellular component GO terms. All annotated human genes were chosen as the statistical domain scope. The significance level was determined using the BH algorithm (FDR <0.05). The Kyoto Encyclopedia of Genes and Genomes (KEGG) database [32] was used to query the most impacted biological pathways.

### Loss-of-function analysis

Tolerance to loss-of-function (LoF) variants was measured using the probability-of-being-LoF-intolerant (pLI) method and gene LoF metrics were obtained from gnomAD (v2.1.1, https://gnomad.broadinstitute.org/) [33]. Genes depleted for null variants were defined as having pLI > 0.9.

### Bootstrapping analysis

Bootstrapping analysis (n=10,000 iterations) was performed to test if observed overlaps were non-random. Each bootstrap iteration generated samples of the same size as in the tested sample for tested condition. The number of shared items (e.g. SNPs) among conditions was counted for each bootstrap iteration. After 10,000 iterations we counted those instances where the number of shared items in the bootstrapped overlap is greater than or equal to the number of shared items in the observed overlap. The *p* value was calculated as the sum of these instances divided by the total number of iterations n,

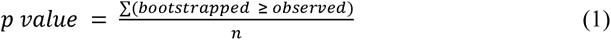

If the *p* < 0.01 we assume that the observed relationship is non-random.

For SNPs analysis, we resampled SNPs from the list of all GWAS SNPs with gwas *p* <5×10^-8^. For gene analysis, we resampled eGenes from the list of all genes in the genome (GENCODE transcript model version 26).

### Data and code availability

Python (version 3.6.9), R (version 4.0.2) and RStudio (version 1.2.5033) were used for data analysis and visualisation. All datasets and software used in the analysis are listed in Supplementary Table 1. A Dockerfile (including the CoDeS3D pipeline and downstream analyses), all findings, scripts and reproducibility report are available on github at https://github.com/Genome3d/genetic_regulation_in_ASD.

## Results

### ASD-associated SNPs mark putative regulatory regions shared between or specific to adult and fetal cortical tissues

ASD-associated SNPs (*p* < 5×10^−8^, n=454) were downloaded from the GWAS Catalog (Supplementary Table 2). Of 454 ASD-associated SNPs, 344 SNPs were represented in both fetal and adult cortex eQTL databases, and were run through the CoDeS3D pipeline (Figure 1, Supplementary Table 3). We identified 80 eQTLs that are involved in 131 significant spatial eQTL-eGene interactions in fetal cortex; and 58 eQTLs that are associated with 67 significant spatial eQTL-eGene interactions in adult cortex (Figure 2a, Supplementary Table 3).

**Fig. 2.**
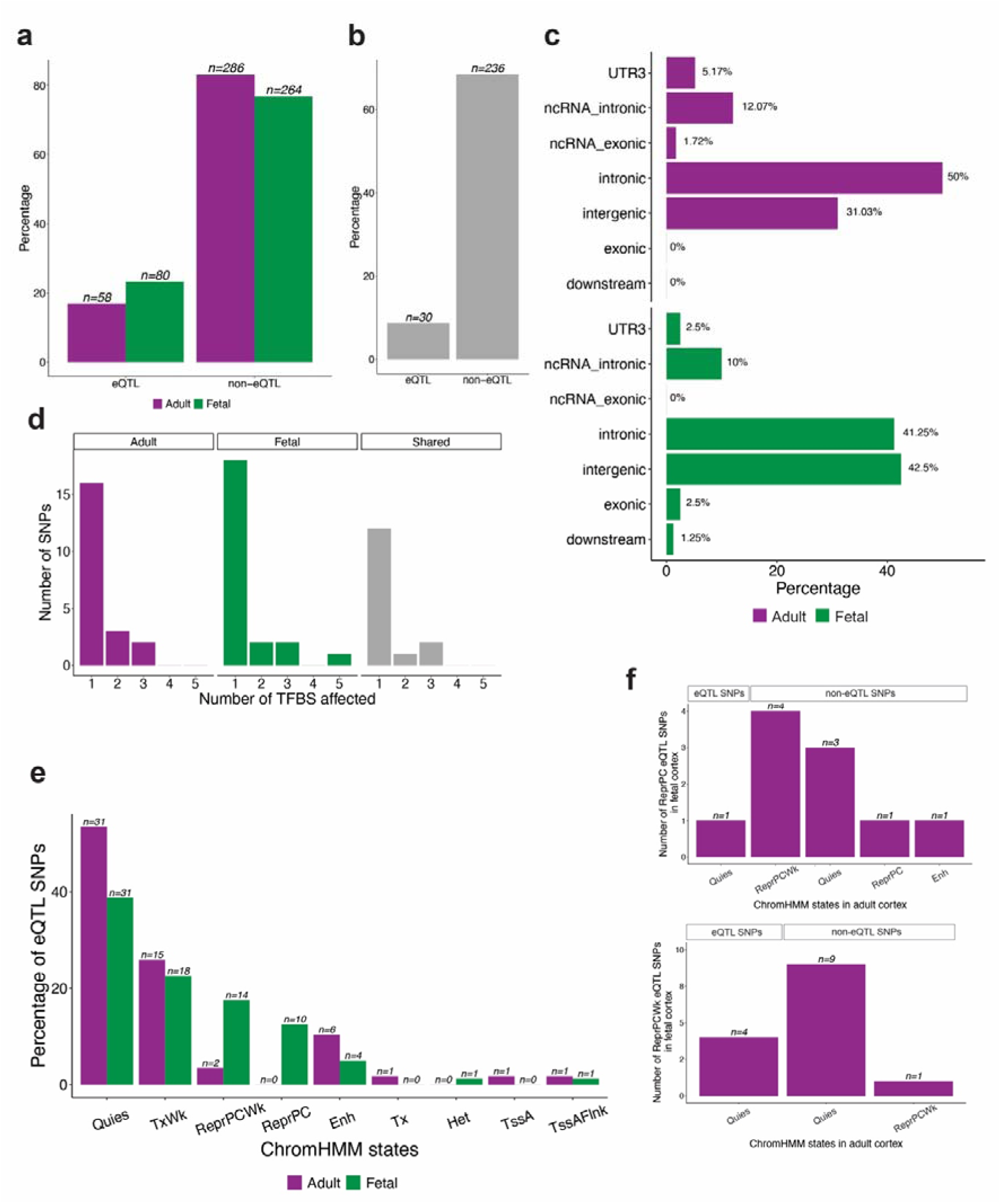
ASD-associated SNPs are enriched within non-coding putative regulatory regions. **a.** Of 344 ASD-associated SNPs represented in both fetal and adult cortex-specific eQTL databases, more SNPs (n=80) are involved in spatial eQTL-gene interactions in the fetal cortex than in the adult cortex (n=58). The proportions of eQTL and non-eQTL SNPs are significantly different in fetal and adult cortical tissues (Fisher’s exact test, *p* = 0.04531). **b.** Thirty ASD-associated SNPs are eQTLs in both fetal and adult cortical tissues. **c.** All ASD-associated eQTLs in adult cortex (n=58) and approximately 78 (97.5%) of the ASD-associated eQTLs within the fetal cortex are located within non-coding genomic regions (Supplementary Table 5). **d**. 15 and 18 ASD-associated eQTLs affect at least one transcription factor binding sites within the fetal and adult cortical tissues, respectively. **e.** Most of the fetal ASD-associated eQTLs are located within quiescent/low transcribed, week repressed PolyComb, repressed PolyComb and weak transcription regions. By contrast, the majority of ASD-associated eQTLs, that were identified in the adult cortex, are located within quiescent/low transcribed and weak transcription regions. Enh=Enhancers, Het=Heterochromatin, Quies=Quiescent/Low, ReprPC=Repressed PolyComb, ReprPCWk=Week Repressed PolyComb, TssA=Active TSS, TssAFlnk=Flanking active TSS, Tx=Strong transcription, TxWk=Weak transcription. **f.** The majority of the fetal ASD-associated eQTLs that are located within weakly repressed PolyComb (ReprPCWk) and repressed PolyComb (ReprPC) regions were not identified as being eQTLs within the adult cortex.

Of the 80 fetal and 58 adult eQTLs, 30 were observed in both fetal and adult cortical tissues (Figure 2b). Fifteen of these shared eQTLs control the same eGenes in fetal and adult cortex (e.g. rs10791097-*SNX19*, rs11191419-*AS3MT*, rs7085104-*AS3MT*, rs174592-*FADS1*; Supplementary Table 4). However, the remaining 15 eQTLs are associated with different eGenes in adult and fetal cortical tissues (Supplementary Table 4). For example, rs35828350: 1) upregulates *NMB* in fetal cortex. *NMB* encodes the neuromedin B peptide that regulates physiological processes including cell growth, exocrine and endocrine secretion [34]; and 2) downregulates *WDR73* in adult cortex. *WDR73* encodes the WD Repeat-containing protein 73 that is linked to microtubule organization and dynamics. Notably, eQTLs involving rs13218591 and rs2237234 regulate different butyrophilin alleles (i.e. *BTN2A2* and *BTN3A1* in the adult and fetal cortex, respectively). The butyrophilin genes encode proteins that belong to the immunoglobulin superfamily and help modulate the immune system [35]. Finally a number of fetal (n=50) and adult (n=28) cortex-specific eQTLs were observed (Supplementary Table 3). Collectively, these observations are consistent with changes in the regulation of subsets of stable and remodelled spatial eQTLs, over the course of brain development, being associated with a predisposition to ASD.

### ASD-associated eQTLs are linked to onset of psychiatric disorders

Previous research has reported shared neurobiological and cellular processes associated with differences in cortical thickness across six psychiatric disorders (i.e. ASD, attention-deficit/hyperactivity disorder (ADHD), bipolar disorder, unipolar depression, obsessive-compulsive disorder and schizophrenia), implicating common mechanisms underlying cortical development [36]. To evaluate possible commonalities among ASD and other phenotypes at the eQTL level in the fetal and adult cortex, we intersected the identified ASD-associated eQTLs with SNPs associated with other traits in the GWAS catalog (*p* <5×10^-8^, assessed on 26/08/2020). Fetal and adult ASD-associated eQTLs were also associated with schizophrenia, unipolar depression, ADHD, bipolar disorder, anorexia nervosa and obsessive-compulsive disorder (Supplementary Figure 2, Supplementary Table 5). We observed that schizophrenia has the largest significant overlap with ASD-associated eQTLs both in fetal (78 out of 80, bootstrapping *p* < 0.01, n=10,000) and adult (57 out of 58, bootstrapping *p* < 0.01, n=10,000) cortical tissues (Supplementary Figure 2, Supplementary Table 5). This observation is consistent with: 1) a comorbid association between ASD and schizophrenia [37]; or 2) a lack of resolution and precision in defining the ASD and schizophrenia phenotypes, and thus possible false positives in GWASs [38].

### ASD-associated eQTLs are involved in diverse regulatory processes

We functionally annotated the ASD-associated eQTLs to understand the potential regulatory mechanisms of the regions they tagged (Supplementary Table 5). As expected, the majority of identified eQTLs were located within intronic and intergenic regions (Figure 2c). The SNP2TFBS [25] database was queried to identify eQTLs that are predicted to alter the affinity of transcription factor binding sites (TFBSs). We identified 21 and 23 eQTLs that reduce the affinity of at least one TFBS in fetal and adult cortical tissues, respectively (Figure 2d). Finally, we tested for enrichment of ASD-associated eQTLs within active regulatory elements and histone modification marks, using ChromHMM [26] 15-state models for adult dorsolateral prefrontal cortex and fetal brain. Fetal ASD-associated eQTLs were located within quiescent/low transcribed (n=31), weak transcription (n=18), week repressed Polycomb (n=14) and repressed Polycomb (n=10) regions (Figure 2e). By contrast, adult ASD-associated eQTLs were located within quiescent/low transcribed (n=31) and weak transcription (n=15) regions (Figure 2e). There was significant enrichment (*p* < 0.01, permutation test: 1000) of ASD-associated eQTLs within loci repressed by Polycomb proteins in the fetal cortex when compared to the adult cortex. Most of these fetal ASD-associated eQTLs located within the Polycomb-repressed eQTLs were not identified as eQTLs in the adult cortex (Figure 2f). Notably, Polycomb repressive complexes have distinct regulatory roles in identity, proliferation and differentiation of neuronal progenitor cells during development [39, 40].

### ASD-associated eQTLs are developmental stage-specific

There is no fundamental reason why the same regulatory elements must or must not impact on the same gene in different tissues, or at different stages of development. ASD-associated eQTLs regulate 81 genes in fetal and 44 genes in adult cortical tissues (Figure 3). Of these genes, 15 are spatially regulated in both fetal and adult cortical tissues (Figure 3, Supplementary Table 4). Different eQTLs are often associated with the gene transcript levels, although the effects of the minor allele are typically similar for the gene in question (i.e. associated with an increase or reduction in transcript levels; Supplementary Figure 3). For example, rs4647903, rs2535629, rs221902, rs7743252, rs832190 eQTLs were associated with increased transcript levels of *DDHD2*, *ITIH4*, *PCNX1*, *TAP2*, *THOC7* genes both in fetal and adult cortical tissues. However, despite having the same effects in fetal and adult tissues, some eQTLs had opposite direction of effects on the gene of interest (e.g. the effects of rs7432375 and rs7618871 on *PCCB* gene transcript levels are not collinear; Supplementary Figure 3). Moreover, we identified development stage-specific eQTLs that were associated with gene transcript levels in either fetal, or adult cortical tissue (Supplementary Figure 3). For example, rs4481150 eQTL is present in both fetal and adult eQTL databases. However, rs4481150 is only associated with increased transcript levels for *ITIH4* in adult cortex. By contrast, the rs1518367 eQTL is associated with reduced transcript levels for *SF3B1* only in fetal cortex. *HLA-DMA* and *BAG6* gene transcript levels are associated with distinct sets of eQTLs in both fetal and adult cortical tissues (Supplementary Figure 3).

**Fig. 3.**
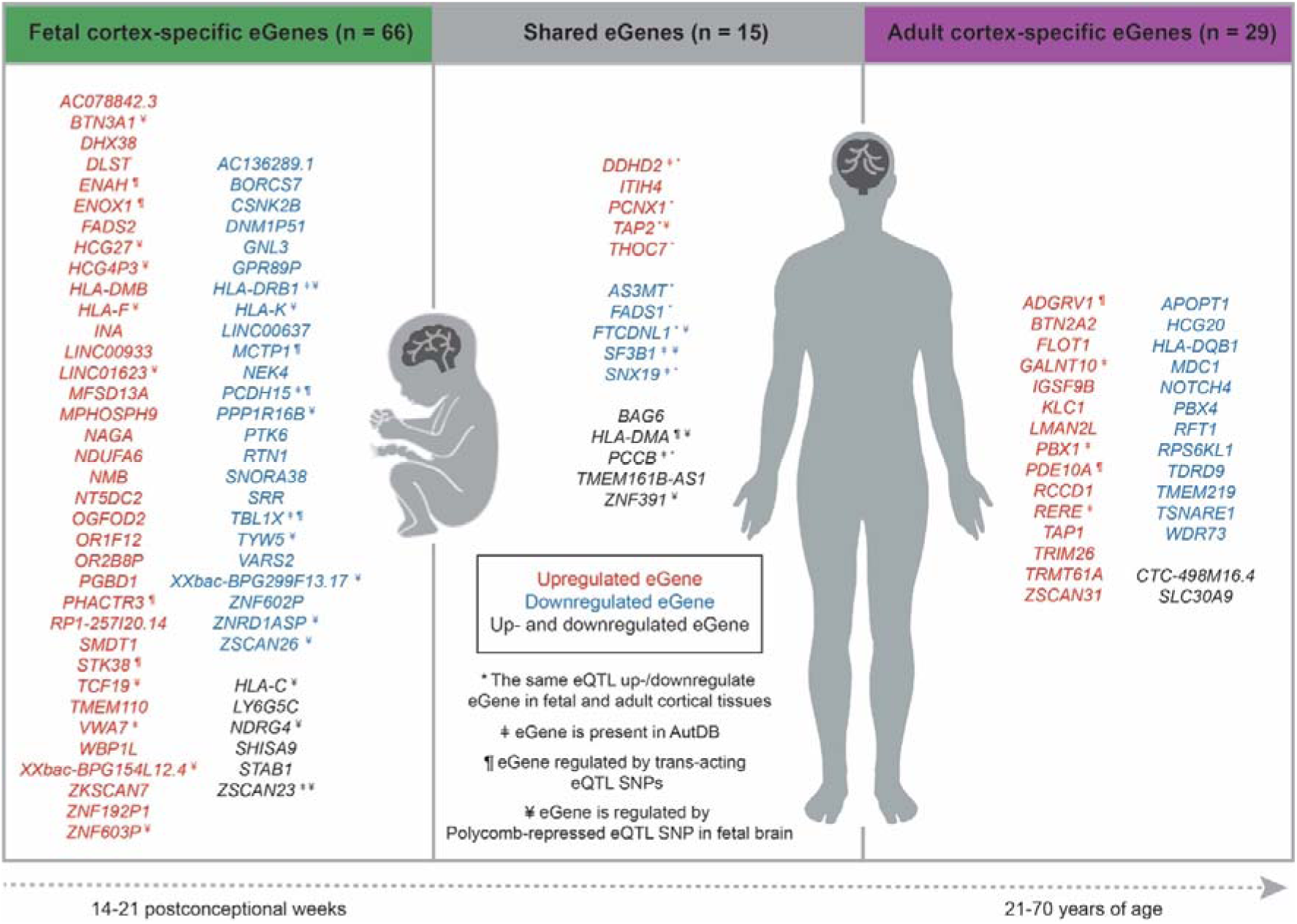
ASD-associated eQTLs mark loci that spatially regulate the expression of genes involved in the fetal brain, in the adult brain, or both. Transcript levels for 15 spatially regulated genes were altered by ASD-associated eQTLs in both the fetal and adult cortical tissues, 66 genes were specific to fetal cortex, and 29 eGenes were specific to the adult cortex. *, the same eQTL is associated with transcription levels for the gene in both the fetal and adult cortical tissues. □, genes that have been curated as being involved in ASD (AutDB [41]; http://autism.mindspec.org/autdb/Welcome.do, assessed on 16/11/2020). ¶, genes whose transcript levels are associated with a trans-acting ASD-associated eQTL. ¥, genes regulated by Polycomb-repressed ASD-associated eQTLs in the fetal cortex.

Despite the existence of fifteen genes in eQTL with ASD-associated SNPs in both fetal and adult cortical tissues, the majority of the changes in gene transcript levels were specific to either fetal (n=66), or adult (n=29) cortical tissue (Figure 3). Transcript levels for these genes are associated with 65 eQTLs in fetal cortex and 39 eQTLs in adult cortex (Supplementary Table 3).

Of the 66 fetal cortex genes, transcript levels for 36 are upregulated in association with a change of the eQTL SNP to the minor allele, while 24 are reduced (Figure 3). Notably, 6 genes are associated with multiple eQTLs which exhibit opposing effects on transcript levels (Figure 3). Nineteen eGenes are regulated by Polycomb-repressed eQTL SNPs, and seven eGenes – by trans-acting eQTL SNPs in fetal cortex (Figure 3).

Similar changes occur in adult cortex where transcript levels for 15 genes are upregulated in association with a change to the minor allele at the eQTL SNP (Figure 3). Again transcript levels for 12 genes are reduced. We also identified 2 genes that had multiple eQTLs where substitution of the SNP with the minor allele had opposing associations with the gene’s transcript levels (Figure 3). Collectively, these findings are consistent with a subset of ASD-associated eQTLs acting in a combinatorial and development stage specific manner to affect the risk of developing ASD.

### Eleven genes associated with ASD-eQTLs have previously been linked to ASD risk

To identify existing and novel gene associations, we intersected our lists of genes, from fetal and adult cortical tissues, with a curated list of 1,237 genes that had been previously implicated in autism development (AutDB [41], accessed on 16/11/2020). Eleven genes (i.e. 8 from fetal cortex: *DDHD2*, *HLA-DRB1*, *PCCB*, *PCDH15*, *SF3B1*, *SNX19*, *TBL1X*, *VWA7*; and 7 from adult cortex: *DDHD2*, *GALNT10*, *PBX1*, *PCCB*, *RERE*, *SF3B1*, *SNX19*) had been previously linked to ASD (Figure 3). Bootstrapping analysis revealed that these overlaps are significant (*p* < 0.01, n=10,000). However, more than 84% of the identified spatially regulated genes were ‘novel’ and have not previously been linked to autism or curated in AutDB.

### ASD-eQTL associated gene set is enriched for loss-of-function tolerant genes

Genes that have essential functions show a decreased tolerance for loss-of-function (LoF) mutations [33]. LoF analysis revealed that 59% (fetal) and 77% (adult) of the eGenes are tolerant to variation that alters the gene sequence. By contrast, 9 fetal cortex-specific genes (i.e. *PHACTR3*, *BAG6*, *CSNK2B*, *SF3B1*, *PPP1R16B*, *FADS2*, *RTN1*, *TBL1X* and *ENAH*) and 5 adult cortex-specific genes (i.e. *PDE10A*, *PBX1*, *SF3B1*, *BAG6* and *RERE*) were LoF intolerant (Supplementary Table 6).

### ASD-eQTL associated genes are enriched for immune-related processes

Functional gene ontology enrichment analysis identified immune-related processes (e.g. antigen processing and presentation) as being enriched in the ASD-eQTL associated gene sets for both fetal and adult cortical tissues (Supplementary Figure 4). Changes to genes within the immune-related processes within adult cortex mostly affect the processing of exogenous antigen. By contrast, immune-related genes that are associated with ASD-eQTLs within the fetal cortex have been implicated in the processing of both endogenous and exogenous antigens (Supplementary Table 7). Removal of all *HLA* genes from the analysis, identified enrichments for genes involved in fatty acid metabolism and processes related to the endoplasmic-reticulum-associated protein degradation (ERAD) pathway within fetal cortex. Removal of *HLA* genes from analyses of the adult cortex gene set identified a retained enrichment for immune-related processes (e.g. antigen processing and presentation), protein kinase C signalling and regulation of cell-cell adhesion processes (Supplementary Table 8).

### ASD-specific protein-protein interaction networks in adult and fetal cortical tissues

The protein-protein interaction (PPIs) network serves as a foundation for cellular signalling circuitry, which mediates cellular responses to environmental and genetic cues. Understanding how ASD-eQTLs affect fetal and adult cortex PPIs could lead to the identification of the pathways that affect cortical development and ASD susceptibility.

PPI data was retrieved from STRING [29] (version 11.0; 24/09/2020). Adult and fetal cortex-specific PPIs (CSPPI) were generated. From these CSPPI networks we identified 42 ASD-associated PPIs within the fetal gene set, and 10 ASD-associated PPIs from the adult cortical tissue gene set (Figure 4, Supplementary Table 9). Louvain clustering analysis identified seven highly connected PPI modules within fetal cortex. KEGG pathway analysis [32] of these modules revealed that they are associated with immune pathways, fatty acid metabolism, aminoacyl-tRNA biosynthesis, spliceosome, ribosome biogenesis in eukaryotes and two modules were not enriched for specific pathway (Figure 4). By contrast, the adult PPI gene set contained three highly connected modules, two of which were not associated with specific pathway, and one of which was enriched for immune pathways.

**Fig.4.**
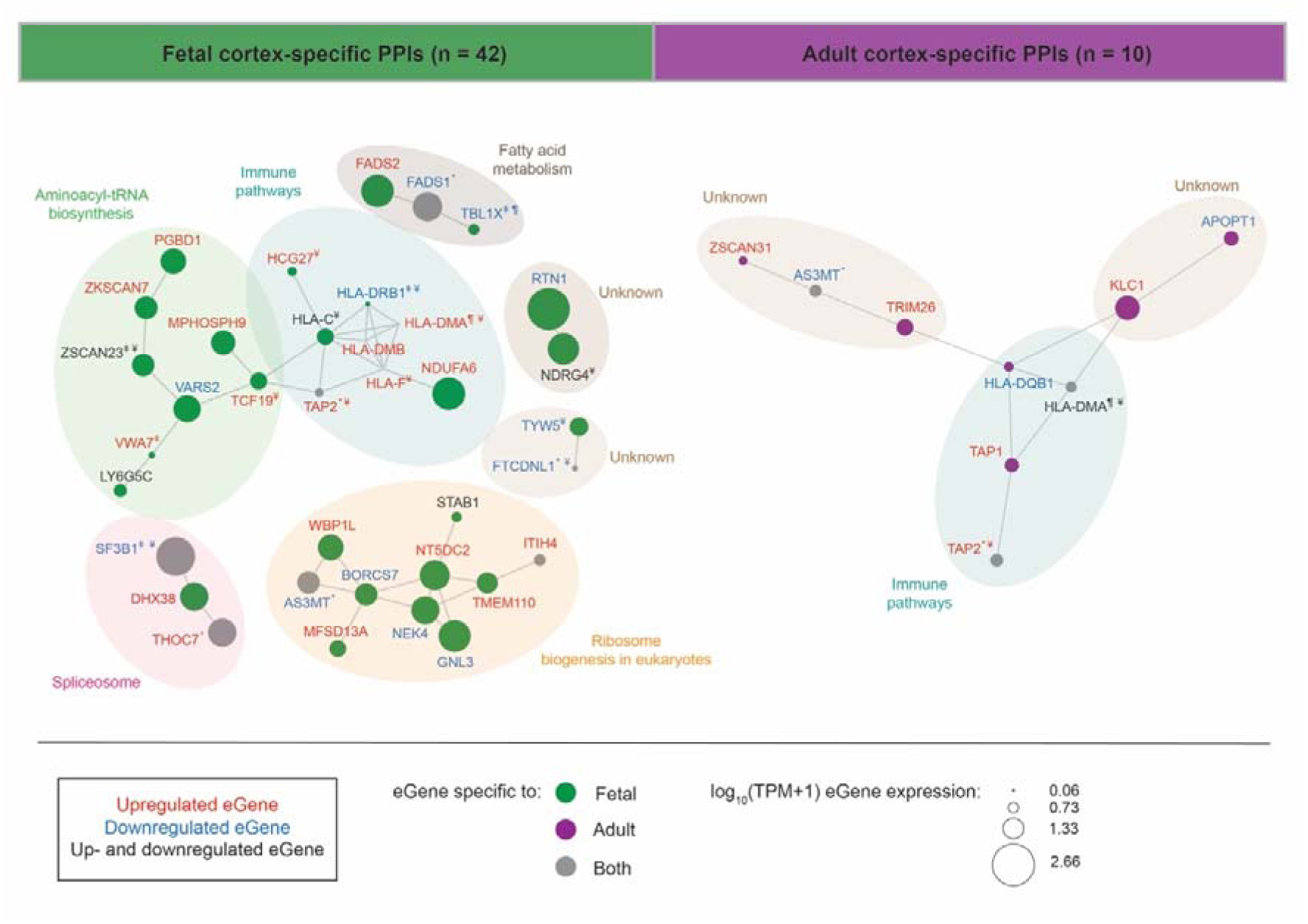
Fetal and adult cortical tissue-specific PPI networks with immune and growth related phenotypes are affected by ASD-associated eQTLs. We identified 42 PPIs in the fetal and 10 PPIs in the adult cortical tissues. Louvain clustering identified seven PPI modules within the fetal cortex that were enriched for immune pathways, fatty acid metabolism, aminoacyl-tRNA biosynthesis, spliceosome, ribosome biogenesis in eukaryotes and two unknown modules. Adult PPIs form three modules that were enriched in immune pathways and two unknown modules. *, gene transcript levels are associated with the same eQTL in both the fetal and adult cortical tissues. □, genes that have been curated as being involved in ASD (AutDB [41]; http://autism.mindspec.org/autdb/Welcome.do, assessed on 16/11/2020). ¶, genes whose transcript levels are associated with a trans-acting ASD-associated eQTL. ¥, genes regulated by Polycomb-repressed ASD-associated eQTLs in the fetal cortex.

The fetal immune PPI cluster contains both MHC class I (i.e. *HLA-C* and *HLA-F*; associated with endogenous antigen processing) and MCH class II (i.e. *HLA-DRB1*, *HLA-DMB* and *HLA-DMA;* associated with exogenous peptide processing) genes. Five genes within this cluster (i.e. *HCG27*, *TAP2*, *HLA-F*, *HLA-DMA* and *HLA-DMB*) are not highly expressed (TPM< 3) within fetal cortical tissue. However, switching the ASD-eQTL to the minor allele is associated with an increase in transcript levels for these genes within the fetal cortex (Figure 4). By contrast, the rs3129968 minor allele is associated with a reduction in transcript levels for *HLA-DRB1*, which is expressed at low levels (TPM<1.25) within fetal cortex. ASD-eQTLs were also associated with increases in *HCG27* (TPM=3) and *NDUFA6* (TPM=116.31) transcript levels within fetal cortex. Notably, the transcript levels for 6 genes in the fetal immune cluster (*HCG27*, *TAP2*, *HLA-F*, *HLA-DMA*, *HLA-C* and *HLA-DMB*) are associated with regulation by Polycomb-repressed ASD-eQTLs.

ASD-eQTLs within the fetal cortex PPI aminoacyl-tRNA biosynthesis cluster are associated with: a) increases in transcript levels for five genes (i.e. *PGBD1*, *ZKSCAN7*, *MPHOSPH9*, *TCF19* and *VWA7*); b) decreases in transcript levels for *VARS2*; and c) two genes (i.e. *ZSCAN23* and *LY6G5C*) whose transcript levels increase or decrease, dependent upon specific eQTL. Similarly, the “ribosome biogenesis in eukaryotes” cluster contained ASD-eQTL associated genes whose transcripts were increased (n=5), decreased (n=4), and one that was subject to increases or decreases in transcript levels depending on specific ASD-eQTL. By contrast, the ASD-eQTL associations within the fetal cortex PPI fatty acid metabolism and spliceosome clusters are less complex (decreases: *FADS1*, *TBL1X, SF3B1,* or increases: *FADS2, DHX38* and *THOC7*; Figure 4). *SF3B1* and *THOC7* were also associated with ASD-eQTLs within the adult cortex. The PPI clusters with unknown functions (Figure 4), contained genes (i.e. *TYW5*, *FTCDNL1*, and *RTN1*) whose transcript levels decrease with the ASD-eQTL and *NDRG4*, whose transcript levels increase or decrease dependent upon the ASD-eQTL.

The patterns of transcript changes in the adult cortex PPI network were similar to those observed in the fetal cortex. Transcript levels for four genes: a) increased (i.e. *TAP1* and *TAP2*); b) decreased (*HLA-DQB1*); or c) both increased and decreased depending on specific ASD-eQTLs (*HLA-DMA*; Figure 4). Both *HLA-DQB1* and *HLA-DMA* are examples of the MCH class II genes that are associated with processing of exogenous antigen. The transcript levels of the genes within the two unknown adult cortex PPI clusters increased (i.e. *KLC1, ZSCAN31* and *TRIM26*), or decreased (i.e. *APOPT1, AS3MT*) with the ASD-eQTL.

## Discussion

In this study, we integrated four distinct levels of biological information (GWAS, eQTL, genome organization [Hi-C] and protein-protein interactions [PPI] networks) to translate genetic variation associated with ASD to the biological pathways that are affected – through alterations to the transcription levels of their component proteins in fetal and adult cortical tissues. We identified shared and development-specific changes to transcript levels for spatially regulated genes within immune pathways. Interestingly, most of the genes within immune-related pathways in fetal cortex are associated with Polycomb-repressed ASD-eQTLs. At the same time, ASD-eQTLs are also associated with regulation of fatty acid metabolism, ribosome biogenesis, aminoacyl-tRNA biosynthesis and spliceosome pathways in fetal cortex. Notably, the significant difference (*p* = 0.04531) in numbers of fetal cortical eQTLs, when compared to adult cortical eQTLs is consistent with a developmental origin for ASD risk. Our findings highlight potential mechanisms through which ASD-associated variants potentially contribute to ASD development (fetal) and onset/maintenance (adult).

We identified ASD-associated eQTLs that mark putative regulatory regions in fetal (n=80) and/or adult (n=80) cortical tissues. It was expected that we would not identify eQTLs for all 344 of the tested ASD-associated SNPs. There are several reasons for this: 1) there are multiple potential mechanisms through which a genetic variant can impact on a phenotype . These mechanisms are not limited to impacts on gene regulation and can affect the splicing activity (so called sQTL SNPs [42]), or trans-acting factors (e.g. non-coding RNAs); 2) ASD is a spectrum disorder not a single highly characterised phenotype; and 3) like all polygenic disorders, ASD is likely to be a whole of body disorder with important etiological contributions from discrete and distant organs within the body.

Studies have previously reported associations between ASD and: 1) schizophrenia [43]; 2) depression [44]; 3) ADHD [45, 46]; 4) bipolar disorder [47]; and other co- and multimorbidities [36, 48, 49]. Our finding that a subset of the identified ASD-associated eQTLs were linked to psychiatric phenotypes highlights the existence of potential shared regulatory mechanisms contributing to the risk of developing these multimorbid conditions. An alternative explanation is that the existence of the shared eQTLs between the multimorbid conditions is due to ambiguity in the phenotyping that was used in the GWAS studies that characterised the phenotype associated-SNPs. However, we contend that these results are consistent with the growing evidence that the vertical approach to connecting genetic variation to phenotype does not adequately account for the multimorbid nature of conditions within the typical variation that is present in humans. Future analyses that incorporate horizontal analyses of all genetic variants associated with ASD and its high-frequency multimorbid conditions will improve our ability to stratify autistic individuals and manage their complications.

Polycomb proteins are known to be involved in transcriptional silencing [50, 51]. However, studies have shown that Polycomb repressive complexes (PRC) can have a dual role in gene regulation during development [52, 53]. For example, PRC1.5 is recruited to activate genes [54] and in combination with AUTS2 (autism susceptibility candidate 2) activates gene expression in neurons [54]. Our results support a dual role for Polycomb – as both a repressor and enhancer of transcription - in the development of ASD risk. Firstly, there was a significant enrichment of ASD-eQTLs within loci that are annotated as being regulated by PolyComb within the fetal, but not adult, cortex. Secondly, the finding that these ASD-eQTLs are associated with both increases and decreases in transcript levels is consistent with the up- and downregulation of the target genes. We contend that empirical studies are required to a) confirm the regulatory activity of the sites (e.g. enhancer reporter assays); b) confirm that the Polycomb complexes are responsible for the observed activity (e.g. by chromatin immunoprecipitation); and c) identify the ‘Polycomb’ subunits that differentiate those sites that enhance or repress transcription within the developing cortex. The results of these experiments would be valuable in identifying novel therapeutic approaches to reduce the risk of full-blown ASD development, particularly given the strength of the evidence for Polycomb roles transcription control [53] and increasing evidence for links to neuronal development (e.g. reviewed in [39]).

Dysregulation of fatty acid metabolism in early brain development may be a risk factor or marker for early-onset of ASD [55]. Abnormalities in lipid metabolism may affect the proper functioning of the nervous system and, thus can contribute to ASD etiology [55–57]. Consistent with this, we identified that ASD-associated genetic variants impact transcript levels for genes involved in fatty acid metabolism in the developing fetal cortex (14-21 postconceptional weeks). Notably, transcript levels for genes within this pathway were not significantly affected by ASD-eQTLs within adult cortex tissues (21-70 years of age). Therefore, with appropriate pre-natal genetic diagnosis of risk and patient stratification, it remains possible that targeted lipid supplementation could reduce the risk of ASD. However, this would require randomised control trials in animals prior to testing in humans. Randomised controlled trials involving pre- and post-conception interventions with different lipids are currently being undertaken or followed up (e.g. [58]) – opening the possibility of post-hoc analyses for ASD risk.

Gene expression is the outcome of numerous processes including transcription, co-transcriptional splicing, mRNA export, and translation. We identified changes within multiple key component pathways of gene expression (i.e. spliceosome and splicing, aminoacyl-tRNA biosynthesis, and ribosome biogenesis) in the fetal cortex. Collectively these results could indicate the existence of a window of tolerable variation within gene expression – outside of which there is risk of developing ASD through changes in global gene expression. Roles for these component gene expression processes in ASD are supported by existing studies (e.g. aberrant splicing and ASD [59, 60]; upregulation of ribosomal protein genes and a higher ribosomal gene dosage can be linked to ASD risk and severity [61, 62]). Moreover, Tărlungeanu et al. (2016) identified a form of ASD resulting from a failure of the brain to properly import amino acids – a critical substrate in gene expression [63]. The significance of these associations in the developing cortex and how they contribute to the etiology of ASD remains to be empirically determined.

Roles for immune dysfunction in ASD etiology are increasingly being identified [64–66]. We also found that immune pathways were affected both in fetal and adult cortical tissues. As we observed for other pathways, there was a developmental separation in the immune pathways that were affected in the fetal and adult cortical tissues. Notably, we found that ASD-eQTL associated changes in transcript levels occurred in adult cortex immune pathways that were associated with processing of exogenous antigen. By contrast, in the fetal cortex changes in gene transcript levels occurred in immune pathways that were related to the processing of both endogenous and exogenous antigens. These findings indicate that there may be developmental stage-specific differences in the impact of the immune system on ASD risk and on-going severity. The existence of changes within fetal cortex pathways for endogenous antigens (e.g. viral) lends some support the hypothesis that the risk of ASD may be greater among children whose mothers suffered from infectious or immune-related diseases during pregnancy - when the infant brain is developing.

The results of our study should be interpreted in view of its strengths and limitations. The main strength of this study is the integration of independent data sets, across distinct biological levels, that include cortex-specific 3D genome structure, eQTL and PPI data with ASD-associated GWAS SNPs. Integrating datasets across biological levels enables us to predict how genetic variation impacts on biological pathways and their involvement in the etiology of ASD. However, our study also has several limitations. Firstly, there is a great phenotypic heterogeneity within autism spectrum disorders, which has led to question their classification into a single diagnostic category [67]. Secondly, common SNPs only account for ∼20% to the ASD risk [6], suggesting that other genetic (e.g., rare variants, structural variation) and environmental factors also contribute to ASD etiology [68, 69]. Increasing the number and sample sizes of the ASD GWAS studies will identify additional genetic variants which may help explain some of this missing heritability [70]. Thirdly, the brain regions that are involved in the etiology of ASD remain difficult to determine. Here, we focused on roles for changes within cortical tissue. However, it is likely that additional regions of the brain (e.g. cerebellum [71]) or other organs are important in the pathophysiology of ASD. As eQTL patterns are tissue-specific, we are unable to extrapolate the results of this study to these tissues. Fourthly, the human brain takes over two decades to build via precisely regulated cell type-specific molecular processes governed by both genetic blueprint and environmental factors. In our study eQTL data represent composite datasets across critical periods of development (e.g. fetal samples were aged from 14-21 postconceptional weeks and the adult samples were from individuals aged 21-70 years). As such, our results only represent snap-shots within the plastic neurodevelopmental trajectory [72, 73]. This is further complicated by tissue averaging across the complex cellular organization and composition which is also different in early development and adulthood. Fifthly, we are aware that the tools and datasets used in this study are potentially biased. For example, identical samples were not used in the ChromHMM, eQTL and Hi-C analyses of the fetal and adult cortical tissues. Furthermore, the Hi-C dataset used to inform the adult cortex analysis consisted of one sample (with one replicate), while two samples (each with three replicates) were used for the fetal cortex analysis (Supplementary Table 1). Finally, protein identifiers (STRING) [29] and transcript identifiers (GTEx and Walker et al. RNA-seq data) [20, 21] were mapped to gene identifiers, thus there was a potential loss of data specificity, since genes typically produce multiple transcripts and protein variants due to alternative splicing.

In conclusion, we have identified clinically relevant putative functional impacts for ASD-associated genetic variants within fetal and adult cortical tissues. We have shown that the transcript levels of genes, whose encoded proteins are known to contribute to immune pathways, fatty acid metabolism, ribosome biogenesis, aminoacyl-tRNA biosynthesis and spliceosome are affected in the fetal cortex. In the adult cortex, the known functions of the impacted genes were enriched in immune pathways. Furthermore, despite not having discussed them in detail, there are number of genes whose transcript levels are affected by ASD-eQTLs whose functions were not enriched within known pathways. Future studies of the roles of these genes in ASD will be important for understanding the full impact of ASD-associated genetic variation in the cortex. We contend that our approach represents a valuable strategy to identify potential ASD candidate genes. Moreover, this approach is not tissue or disease specific and is capable of identifying previously unknown tissue-specific contributions to ASD etiology and its interactions with multimorbid conditions. Similar approaches, in combination with existing and future clinical studies of ASD will contribute to individualized mechanistic understanding of ASD etiology in early brain development and adulthood.

## Acknowledgments

The authors would like to thank the Genomics and Systems Biology Group (Liggins Institute) for useful discussions. EG is the recipient of a Liggins Ph.D. scholarship and was supported by MBIE Catalyst grant (The New Zealand-Australia LifeCourse Collaboration on Genes, Environment, Nutrition and Obesity (GENO); UOAX1611). This work was funded by a University of Auckland FRDF grant (Confirming spatial connections to unravel how SNPs affect phenotype; 3714499) and a MBIE Catalyst grant (The New Zealand-Australia LifeCourse Collaboration on Genes, Environment, Nutrition and Obesity (GENO); UOAX1611) to JOS. JOS was also funded by a Royal Society of New Zealand Marsden Fund [Grant 16-UOO-072]. The Genotype-Tissue Expression (GTEx) Project was supported by the <u>Common Fund</u> of the Office of the Director of the National Institutes of Health, and by NCI, NHGRI, NHLBI, NIDA, NIMH, and NINDS.

## Author contributions

EG performed the analyses and wrote the manuscript. TF contributed to the eQTL analysis and commented on the manuscript. MV and CW co-supervised EG and commented on the manuscript. TL contributed to discussions that aided results interpretation and commented on the manuscript. JOS supervised EG and co-wrote the manuscript.

## Conflict of interest

The authors declare no conflict of interest.

## Supplementary Tables

**Supplementary Table 1:** Datasets and software used in this analysis.

**Supplementary Table 2:** List of ASD-associated SNPs used in this study.

**Supplementary Table 3:** Spatial ASD-associated eQTL - gene interactions in adult and fetal cortex tissues.

**Supplementary Table 4:** ASD-associated eQTLs that are present in both adult and fetal cortex tissues.

**Supplementary Table 5:** Functional annotation of ASD-associated SNPs used in this study.

**Supplementary Table 6:** Loss-of-function analysis results.

**Supplementary Table 7:** Gene Ontology enrichment analysis results.

**Supplementary Table 8:** Biological processes for fetal and adult cortex-specific genes associated with the ASD-associated eQTLs, after removing all *HLA* genes from the lists.

**Supplementary Table 9:** ASD-associated PPIs in fetal and adult cortical tissues.

---

**Supplementary Fig. 1.**
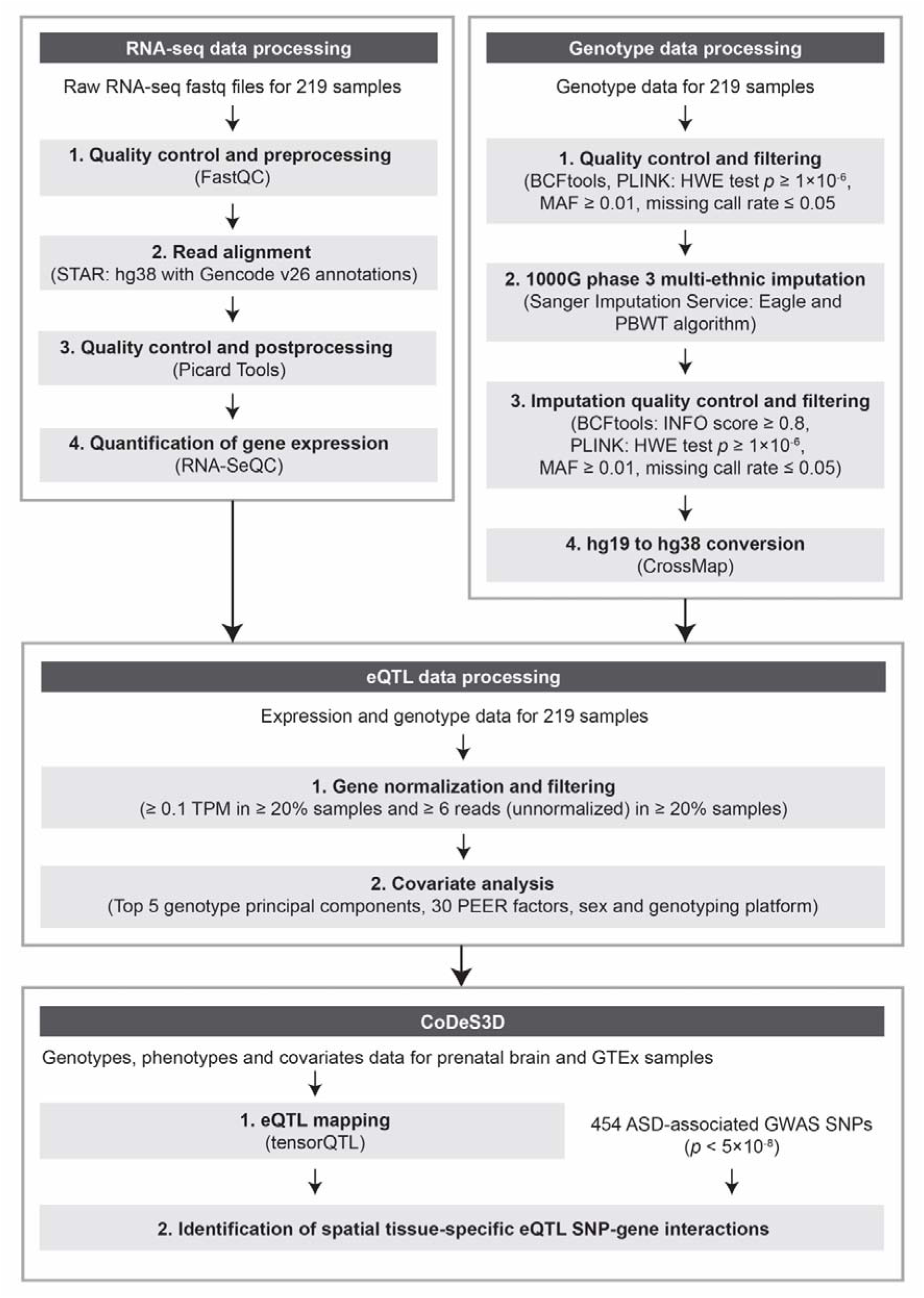
Overview of the pipeline that was used for fetal brain RNA sequencing and genotype data processing. Output data from this pipeline were used as inputs for eQTL analysis and identification of spatial eQTL-gene interactions in the fetal brain, using the CoDeS3D algorithm (Fig. 1).

**Supplementary Fig. 2.**
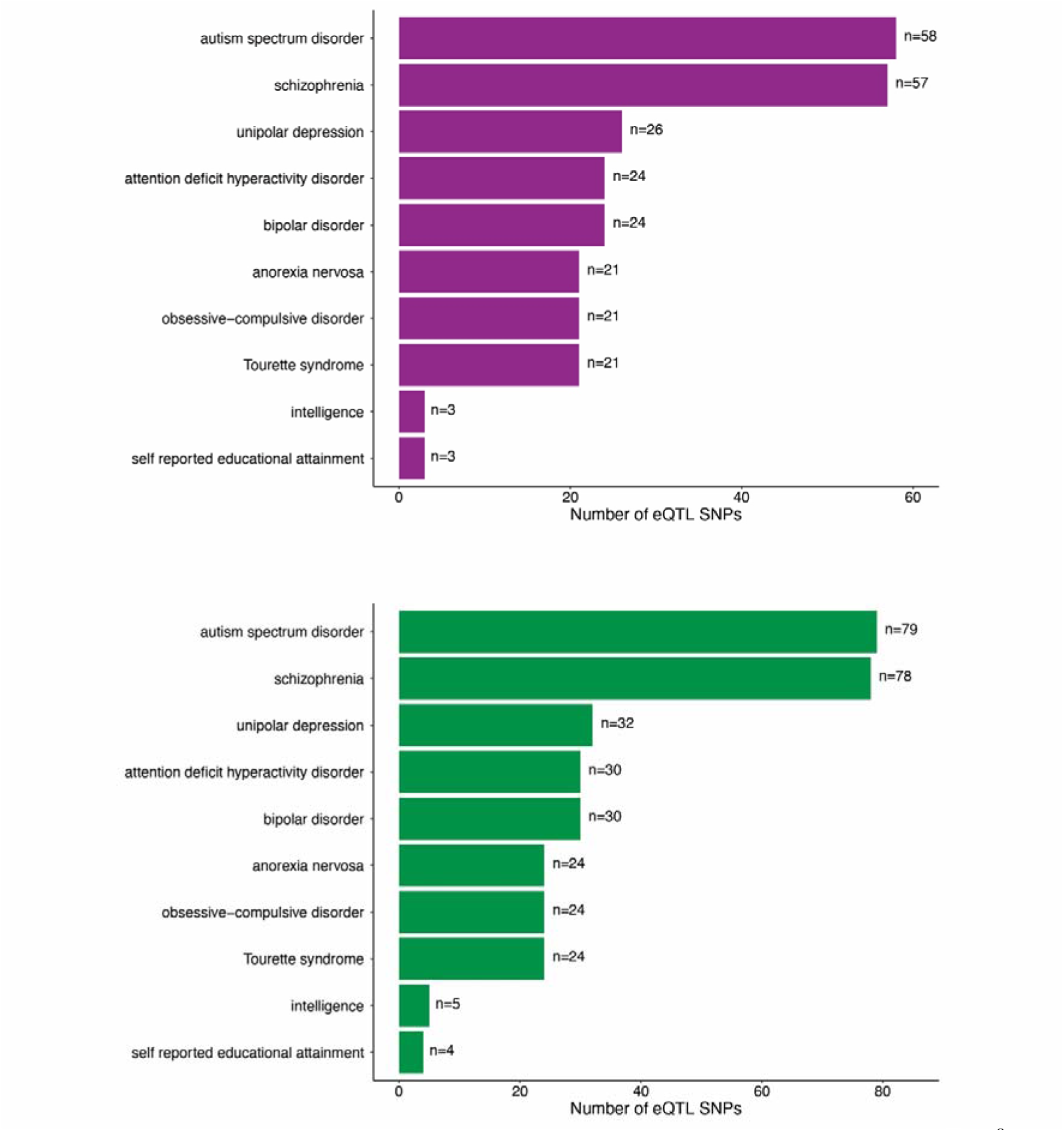
Rank order representation of the top 10 GWAS traits (*p* < 5×10^-8^, assessed on 26/08/2020) for which the ASD-associated eQTLs are annotated as being risk loci. 57 out of 58 ASD-associated eQTL SNPs in adult cortex and 78 out of 79 eQTL SNPs in fetal cortex were associated with schizophrenia. These overlaps are statistically significant (bootstrapping, *p* < 0.01, n=10,000).

**Supplementary Fig. 3.**
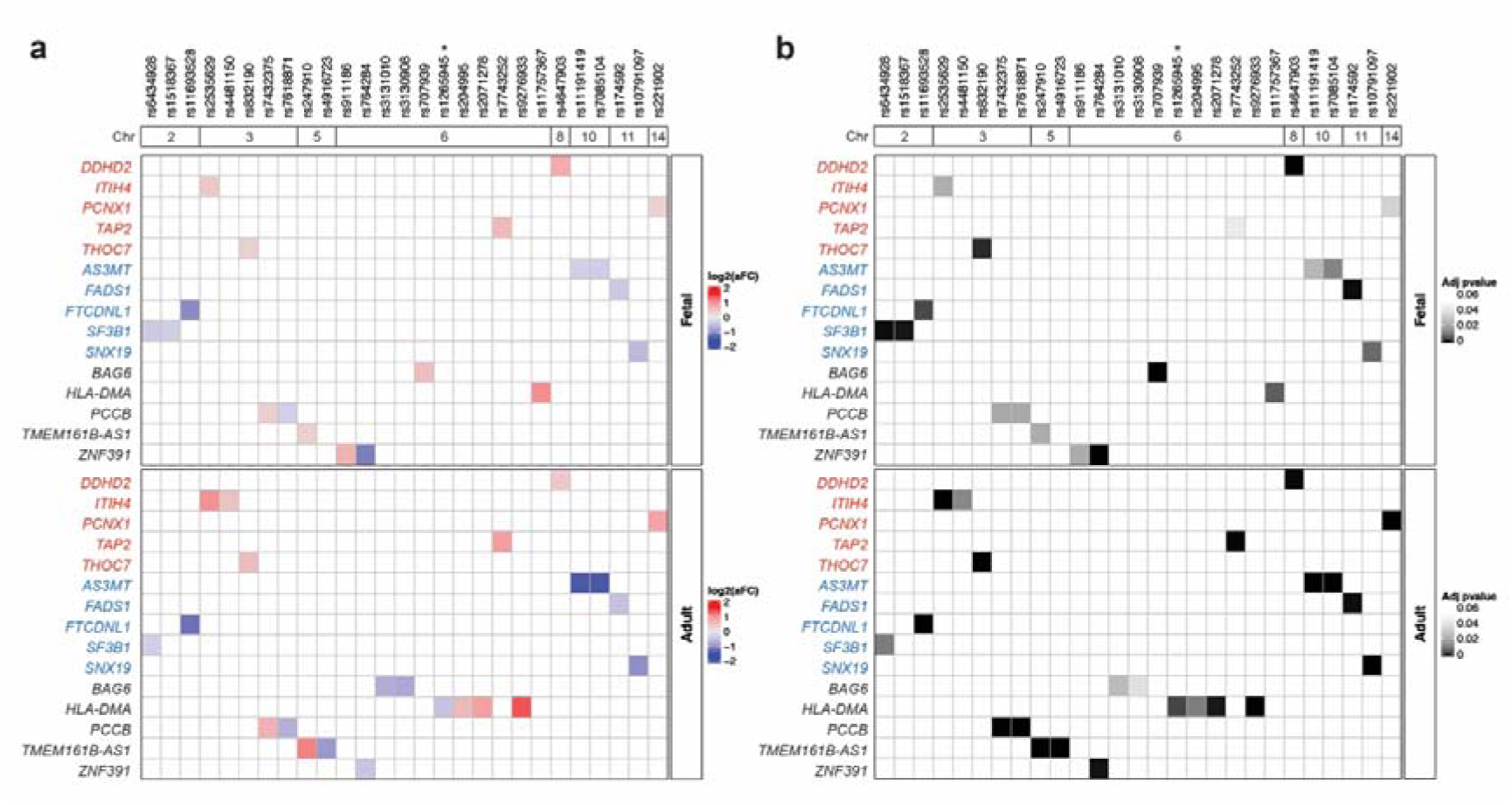
The ASD-associated eQTLs typically affected gene transcript levels collinearly up- and downregulate the expression of 15 eGenes common in both adult and fetal cortical tissues. **a.** ASD-associated eQTLs that present in both tissues typically affect gene transcript levels colinearly. However, specific eQTLs that act only in fetal or adult cortical tissue. **b.** Adjusted *p* (adj *p* < 0.05) for the ASD-associated eQTLs in a. *, gene whose transcript levels were associated with a trans-acting ASD-associated eQTL.

**Supplementary Fig. 4.**
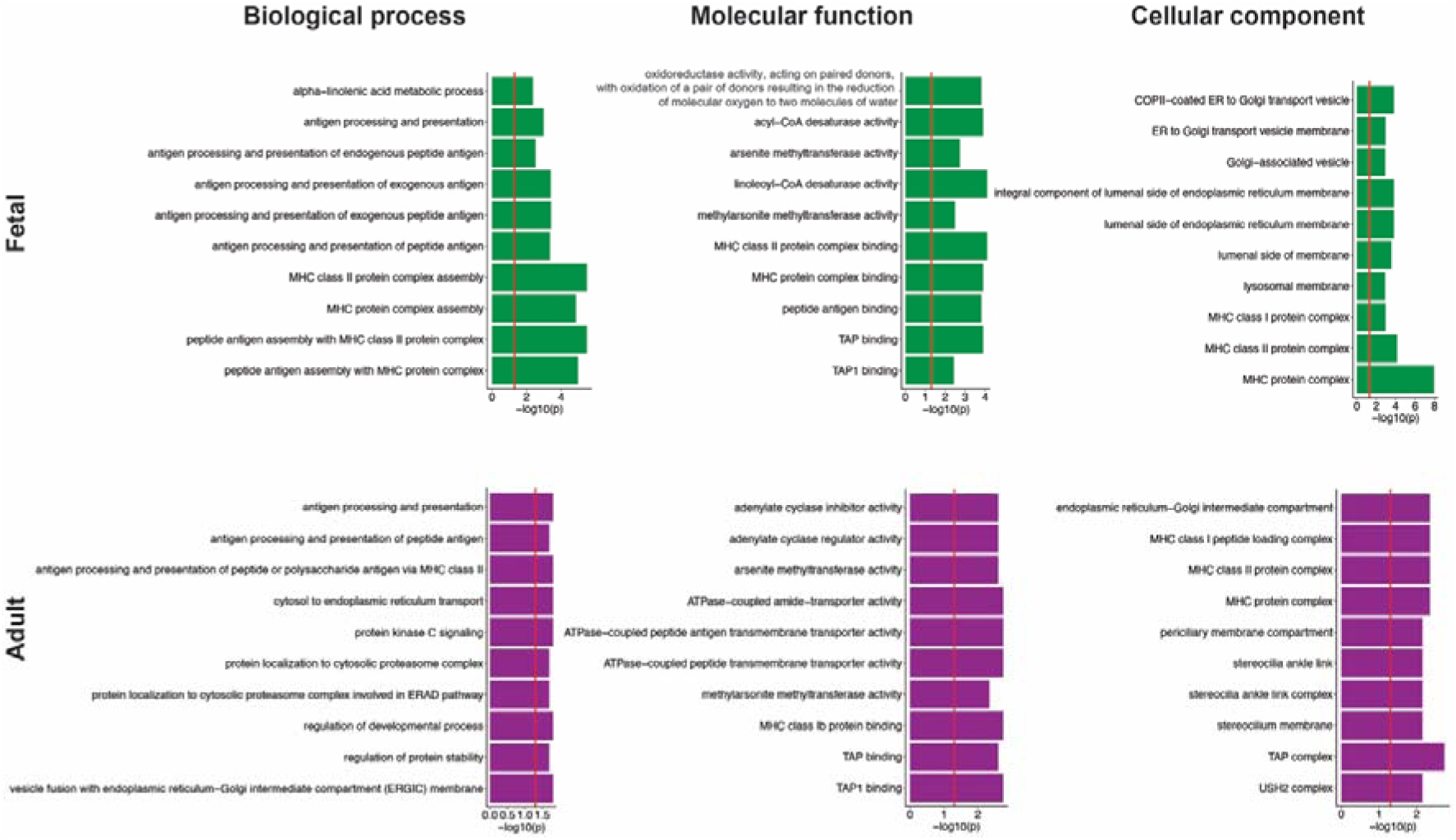
Top 10 biological processes, molecular functions and cellular components for fetal and adult cortex-specific genes associated with the ASD-associated eQTLs. Gene ontology (GO) enrichment analysis was performed using g:Profiler. The threshold for significance (red line) is *p* < 0.05.

## Notes

### Competing Interest Statement

The authors have declared no competing interest.

